# Single-Cell Gene Expression Profiling and Cell State Dynamics: Collecting Data, Correlating Data Points and Connecting the Dots

**DOI:** 10.1101/044743

**Authors:** Carsten Marr, Joseph X. Zhou, Sui Huang

## Abstract

Single-cell analyses of transcript and protein expression profiles – more precisely, single-cell resolution analysis of molecular profiles of cell populations – have now entered the center stage with widespread applications of single-cell qPCR, single-cell RNA-Seq and CyTOF. These high-dimensional population snapshot techniques are complemented by low-dimensional time-resolved, microscopy-based monitoring methods. Both fronts of advance have exposed a rich heterogeneity of cell states within uniform cell populations in many biological contexts, producing a new kind of data that has stimulated a series of computational analysis methods for data visualization, dimensionality reduction, and cluster (subpopulation) identification. The next step is now to go beyond collecting data and correlating data points: to connect the dots, that is, to understand what actually underlies the identified data patterns. This entails interpreting the “clouds of points” in state space as a manifestation of the underlying molecular regulatory network. In that way control of cell state dynamics can be formalized as a quasi-potential landscape, as first proposed by Waddington. We summarize key methods of data acquisition and computational analysis and explain the principles that link the single-cell resolution measurements to dynamical systems theory.

## Introduction

A cell state transition is an elementary event in metazoan development. The associated phenotypic change, e.g. cell differentiation, cell growth termination or artificial cell reprogramming, has traditionally been explained by molecular signaling pathways. Nowadays, a cell state is characterized exhaustively by its molecular profile, e.g. its transcriptome or proteome. However, the characterization of static molecular profiles cannot explain essential properties of the cell state *dynamics*, such as discreteness of states, stability of states, the binary nature of cell transitions and the directionality of cell development. These properties emerge from nonlinear dynamics of the molecular regulatory networks involved, such as the gene regulatory network, that govern cell state dynamics.

Only in the past decade has nonlinear dynamical systems theory entered the center stage in the study of cell state transitions with the renaissance of Waddington’s epigenetic landscape [1] as a conceptual aid in stem cell and developmental biology. Waddington proposed a “landscape” to explain that cells differentiate into discrete, robust cell states. In his view, a marble, representing a cell, rolls down a hilly landscape towards a number of valleys and must eventually settle in one of them - each representing a particular cell type. This landscape is, as we will see, more than a metaphor but has a mathematical basis.

With the arrival of single-cell technologies we can in principle uncover the topography of the landscape by profiling individual cells in as many positions as possible. Technologies for monitoring single-cell states can be divided into two complementary types: measurement (i) of a large number of variables of a cell state (e.g. abundance of transcripts/proteins) as “snapshot” at a given time point in a large number of cells or (ii) of just a handful of variables continuously observed over time in the same cell and its descendants. The former destroys the cells during measurement, the latter keeps cells alive, allowing for longitudinal monitoring and providing information unique to biological systems, such as dependencies between mother and daughter cells. While the analysis of single developing cells has a over 100 year old history (see [2] for a recent review), what is new is the massively parallel nature and the high-dimensionality: a large number of cells can be analyzed simultaneously for a large number of cellular variables. Thus, novel technologies are less about “single-cell”, but rather allow analysis of entire cell populations with single-cell *resolution*.

So far most analyses of single-cell profiles and longitudinal observations are agnostic of the formalism of dynamical systems that underlies the intuitive picture of the landscape. Although sometimes Waddington’s landscape is invoked, current approaches almost exclusively focus on descriptive computational analyses for data visualization, dimension reduction, or statistical pattern identification (see [3,4] for reviews). But now the time is ripe to move beyond collecting the data and correlating the data points, and to connect the dots: We need to apply the formal theory of dynamical systems that is able to explain the uncovered patterns in single-cell data. In the following, we review recent technological developments (collecting the data) and the computational tools as a first level data organization (correlating the data points) before summarizing the interpretation of data in the light of formal concepts of dynamical systems (connecting the dots).

## Collecting the data: snapshot sampling of single cells

Single-cell techniques, if applied to a sufficiently large number of cells, provide distributions of measured variables for the entire population. Such distributions offer an unprecedented wealth of information about the dynamics of cell states, far beyond cell-cell variability and higher statistical moments.

To appreciate this, one needs to first accept the biological fact that variable distributions even in an isogenic uniform population of cells of the nominally same “cell type” do not just reflect inconsequential (thermal) fluctuations in gene expression, let alone technical (measurement) noise, but also a biologically significant diversity of cellular states with functional consequences. For instance, repeated fluorescent activated cell sorting (FACS) analyses have revealed distinct subpopulation dynamics among mouse embryonic stem cells (mESCs) with respect to expression of the pluripotency factor Nanog [5]. It is also evident that individual cell states are not static but dynamic, exposed by the slow noise-driven re-establishment of a heterogeneous marker distribution in hematopoietic cells from a sorted subpopulation [6]. The main limitation for FACS is that the number of proteins that can be simultaneously analyzed barely exceeds a dozen due to the overlap of optical emission spectra. An advancement is single-cell mass cytometry (CyTOF), where antibodies are tagged with heavy metals and measured via mass spectrometry [7–9]. The sharper discrimination allows for up to 50 proteins to be measured simultaneously in each cell.

Single-cell technologies for measuring transcripts progressed tremendously in recent years. Quantitative reverse transcription polymerase chain reaction (qRT-PCR) technology on nanoliter-scale has been applied to nearly 4,000 cells in different stages of early blood development [10]. However, the number of different mRNAs that can be analyzed in one cell is limited in PCR-based approaches. Profiling whole transcriptomes can be achieved with single-cell mRNA sequencing (RNA-Seq), where the crucial step is unbiased amplification of cDNA before sequencing (see [11–13] for reviews). Use of unique molecular identifiers (UMIs) that bar-code each molecule, not just each transcript species allows a robust quantification by intercepting amplification bias [14]. Cell throughput has been pushed upwards recently (see Figure 1) with the combination of biotechnological methods (e.g. barcoding for multiplexing cells to run in the same sequencing reaction, or the use of microfluidics and droplets for initial reactions of individual cells). Using Drop-seq, 39 subpopulations of mouse retinal cells have been identified in 40,000 cells [15] and population heterogeneity in nearly 6,000 mESCs has been profiled [16]. A drawback RNA-Seq in single cells is the reduced sensitivity such that only the 10-20% most abundant transcripts can be quantitated [17,18]. Careful experimental design, balancing the tradeoff between number of cells and transcripts sequenced and sequencing depth [17] and appropriate computational post-processing, e.g. to correct for cell-cycle induced heterogeneities [19], are crucial for single-cell transcriptomics.

**Figure 1:**
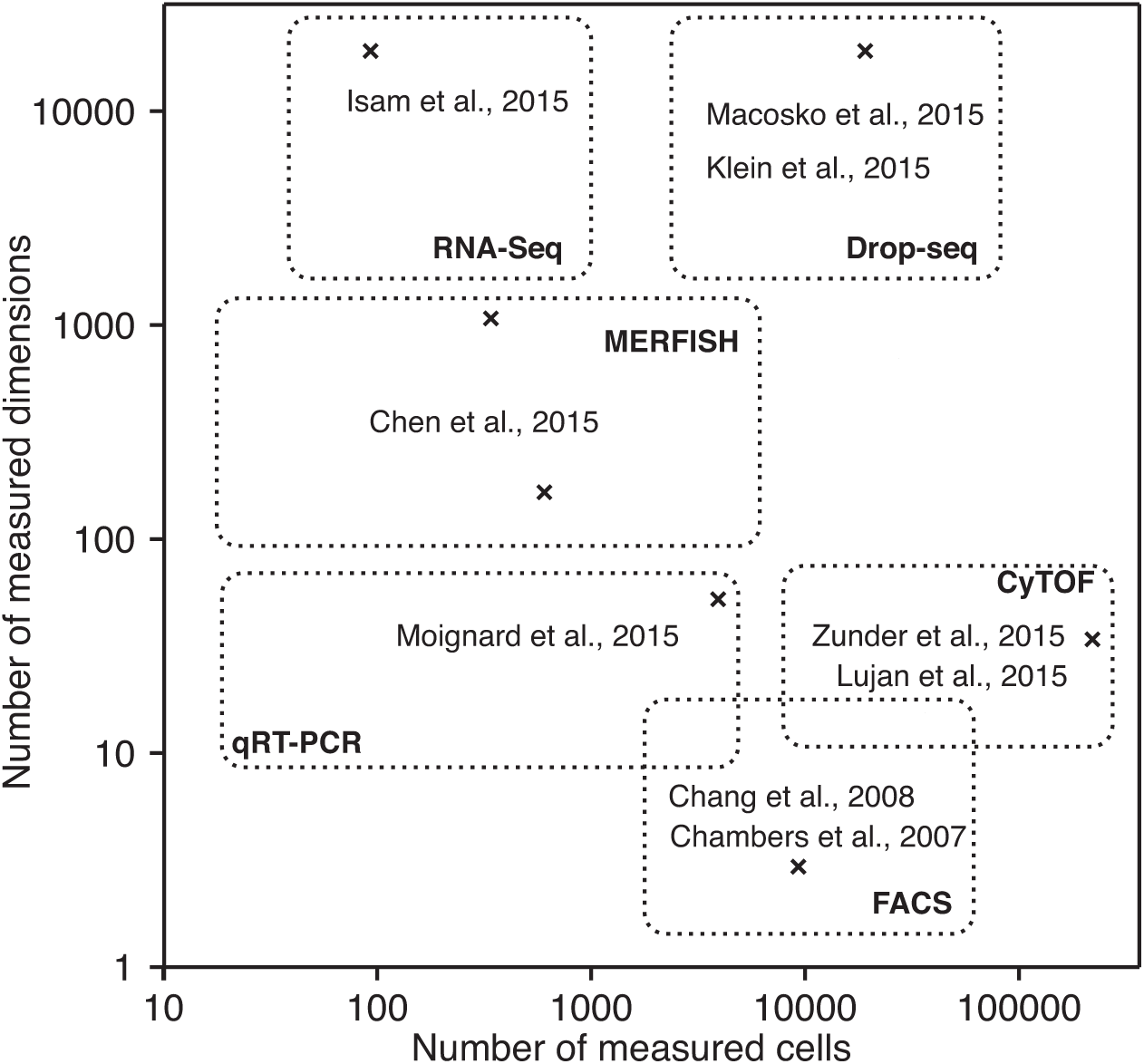
Recent applications of single-cell snapshot technologies sample up to hundreds of thousands of single cells while measuring multiple variables, e.g. genome-wide mRNA expression.

Gene expression manifests cell states only at one particular level. The chromatin state of individual cells can be determined with single-cell bisulfite sequencing for DNA methylation [20], the Hi-C method for chromosome conformation [21], the transposase-accessible chromatin assay (ATAC-seq) [22,23], and immunoprecipitation followed by sequencing (ChIP-seq) for histone methylation [24]. Also the spatial context of single cells is now accessible with recent extensions to multiplexed fluorescent in situ hybridization [25,26], and imaging mass cytometry [27]. Finally, the feasibility of combining genomics and transcriptomics in the same single cell has recently demonstrated [28].

## Correlating the data points: computational data analysis of snapshots

High-throughput (many cells) and high-dimensional (many variables) single-cell measurements provide a wealth of information on molecular profiles in populations. The output for a biological sample is now no longer a vector of *m* components (e.g. the *m different* mRNA species in a single-cell RNA-Seq experiment) as in population/tissue omics, but a [*m*×*n*] matrix since we now measure expression in *n* individual cells. Mathematically, each cell can be positioned in an *m*-dimensional space, where the axes are the measured variables. Using this notation, which is also the basis for the dynamical systems analysis discussed later, the first generation of computational tools has been developed to handle this new type of data: to reduce the m-dimensional space by mapping individual cells onto an interpretable lower (two- or three-) dimensional space with minimal loss of information, to identify patterns, such as an (pseudo)temporal order of cell states or static clusters, and to visualize the data.

Pearson’s principal component analysis (PCA) identifies genes that vary the most within the profiled population of cells and linearly projects the high-dimensional data into a lower dimensional space. It has been used for single-cell transcriptomics to classify mouse sensory neurons into novel cellular subtypes [29]. The incorporation of censoring of expression values due to non-detected transcripts has been achieved by a probabilistic version of PCA [30] and an extension of the factor analysis framework [31]. Nonlinear methods are often better suited to identify the low-dimensional manifolds to which gene expression is confined. One popular method is *t*-SNE – a variant of stochastic neighbor embedding using the Student-t distribution to calculate pointwise similarity [32]. Recently, it has been used to cluster retinal cells [15] and embryonic stem cell populations [16] from droplet-based RNA-Seq. t-SNE preserves local distances and thus ensures that neighboring data points in the original data space are still nearby in the low-dimensional embedding. However, also points distant in the data space could be rather close-by in the embedding. Alternative dimension reduction methods, based on Gaussian processes [33] and diffusion maps [34,35], preserve global state space distances between cells. A comparison of dimension-reduction algorithms for single-cell analyses is provided in [34].

Several approaches have been developed to identify clusters within the high-dimensional data set, a step towards discovering new cell subpopulations of biological significance. Spectral clustering [36] and density-based cell population identification [37] for the analysis of FACS data has been proposed. For CyTOF data, subpopulations have been identified with a regularized regression-based method [38] and a graph-based method with community detection to maximize “modularity” [39]. Cell hierarchies have been estimated based on minimum spanning trees [40], and a divisive bi-clustering method [41] infers classes of molecularly distinct cells in the mouse brain. Of obvious biological interest is also to identify rare cell types outside of abundant clusters. Gruen et al. [42] devised an algorithm to do so and predicted and validated a rare intestinal cell type.

A first hint of awareness of a dynamical process underlying the observed patterns is offered by methods that seek to quantify interrelatedness between cells by viewing them as snapshots of a temporal succession of states despite being measured at the same time. Such “smearing out” of a population is plausible given the stochastic asynchrony of biological processes between cells. The ‘monocle’ package [43] allows the inference of a pseudotemporal order via independent component analysis (ICA) and reconstruction of a mean spanning tree based path through the low dimensional embedding. An alternative algorithm called Wanderlust [44] reconstructs a developmental trajectory based on nearest neighbor graphs in the high-dimensional measurement space. A statistical analysis was used to infer oscillatory genes from a single-cell RNA-Seq snapshot, where unsynchronized cells were mixed [45].

## Longitudinal sampling

The snapshot methods discussed above either destroy the observed sample (e.g. via cell lysis in the case of RNA-Seq and qRT-PCR and laser ablation in the case of imaging mass cytometry), fix cells (in the case of FISH), or lose the identity of individual cells between two measurement points (during repeated FACS). Some biological questions however [46]require monitoring of the molecular state of the very same cell at consecutive time points [46]. This is achieved by traditional video microscopy or, if cytotoxicity is involved and the process of interest is slow, by time-lapse microscopy. Successful implementation requires (i) choice of appropriate markers, (ii) conditions that keep cells in physiological conditions under the microscope and (iii) suitable methods for tracking and analysing individual cells (see [2,47] for reviews).

In mESCs, the dynamics of the heterogeneously expressed pluripotency factor Nanog has been scrutinized in this manner, demonstrating the switching between Nanog-high and Nanog-low cells. Such longitudinal studies quantified gene expression fluctuations across cell cycles, subpopulations, and conditions using reporter systems [48–52] or fusion proteins [53]. To study fundamental aspects of gene expression, mRNA levels have be measured using the MS2 system [49], where a specific target sequence is incorporated in the non-coding portion of the RNA of interest, forming a RNA stem loop that is bound by the constitutively expressed viral MS2 protein fused with a fluorescent protein [54]. Naturally, these live cell imaging methods measure low-dimensional dynamics: they monitor a relatively small number (typically up to two) of state variables, but in turn they provide a wealth of information on the temporal structure of gene expression.

## Connecting the dots: analysis informed by dynamical systems theory

The patterns discovered by descriptive computational analyses, and how they change as cells switch their phenotypic state, must obviously be driven by a force and guided by constraints. A given singlecell state, for instance, can be stable or unstable, fated towards a particular state or capable of choosing between multiple fates [55]. The governing principles of these dynamical properties can be comprehended in a formal manner by considering the regulatory network that controls gene expression. With that we move from the epistemology of phenomenological analysis to that of a formal framework anchored in a set of first principles of physics.

It is in this sense that Waddington‘s landscape enters the interpretation of single-cell data. The landscape manifests the constraints on cell state changes (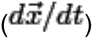)that emanate from the nonlinear dynamics of the underlying regulatory system (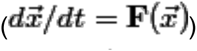) [1,56–58], which is often theoretically embodied by a gene regulatory network (for network inference based on snapshot data see [10,59]). The regulatory interactions between genes generate a driving force on the cell - an analog of the force in Newtonian mechanics. For instance, if gene A encodes an inhibitor of gene B, then as the expression of gene A increases, the expression of gene B will decrease. Such coordinated change of expression across all genes takes place until a cell reaches a stable steady state (where 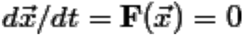 and the system is robust to perturbations), which corresponds to a stable attractor – the point at the bottom of a potential well. Thus, in principle, the entire behavioral repertoire of a cell is encoded in the genome and uniquely maps into a particular landscape [57]. Internal fluctuations of gene expression push the cell away from the attractors – “against the uphill slope”. If the restoring force is not exceeded, the cell will be pushed back to the attractor. For a cell population, this confluence of deterministic and stochastic dynamics gives rise to “clusters” in state space, which account for the non-genetic cell-cell variability. Thus, the spread of the cluster around the attractor state is a measure of heterogeneity of this specific cell type [60]. Being different from a linear dynamical system that can have one attractor at most, nonlinearities in the rate equations describing the dynamics of the network are able to produce multiple solutions – multiple attractor states [61].

In the modern version of Waddington’s landscape [62] we are interested in the relative stability of each attractor – the relative “depth” of the valleys. In this perspective, different cell types have distinct quasi-potentials. A phenotype change induced by an external perturbation can be imagined as an equivalent of “catalysis”: the barrier (hill) between attractors is lowered because of changes in the regulatory interactions conferred by signal transduction. This allows cells to swarm out of the original, now flattening attractor and reach new nearby attractor states [63] (see Figure 2). Thus, even from the qualitative landscape image, the relative depth of an attractor governs the direction of likely transitions. The landscape’s slope embodies the driving force of cell differentiation and the arrow of time of development [64]. In theory, a landscape can be computed numerically, in which the quasipotential 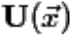(the elevation) reflects the probability of transitions between attractors along a “least effort path”; but this would require knowledge of the system specifications 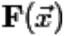 from the governing rate equations 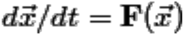 of the dynamical system, i.e., the architecture of the network and reaction modalities of every regulatory interaction. Since such detailed knowledge is not available and constructing 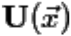 would be computationally expensive if we had the full information, only partial landscapes can be derived from models of known gene-gene interactions, which typically consist of small circuits [65].

**Figure 2:**
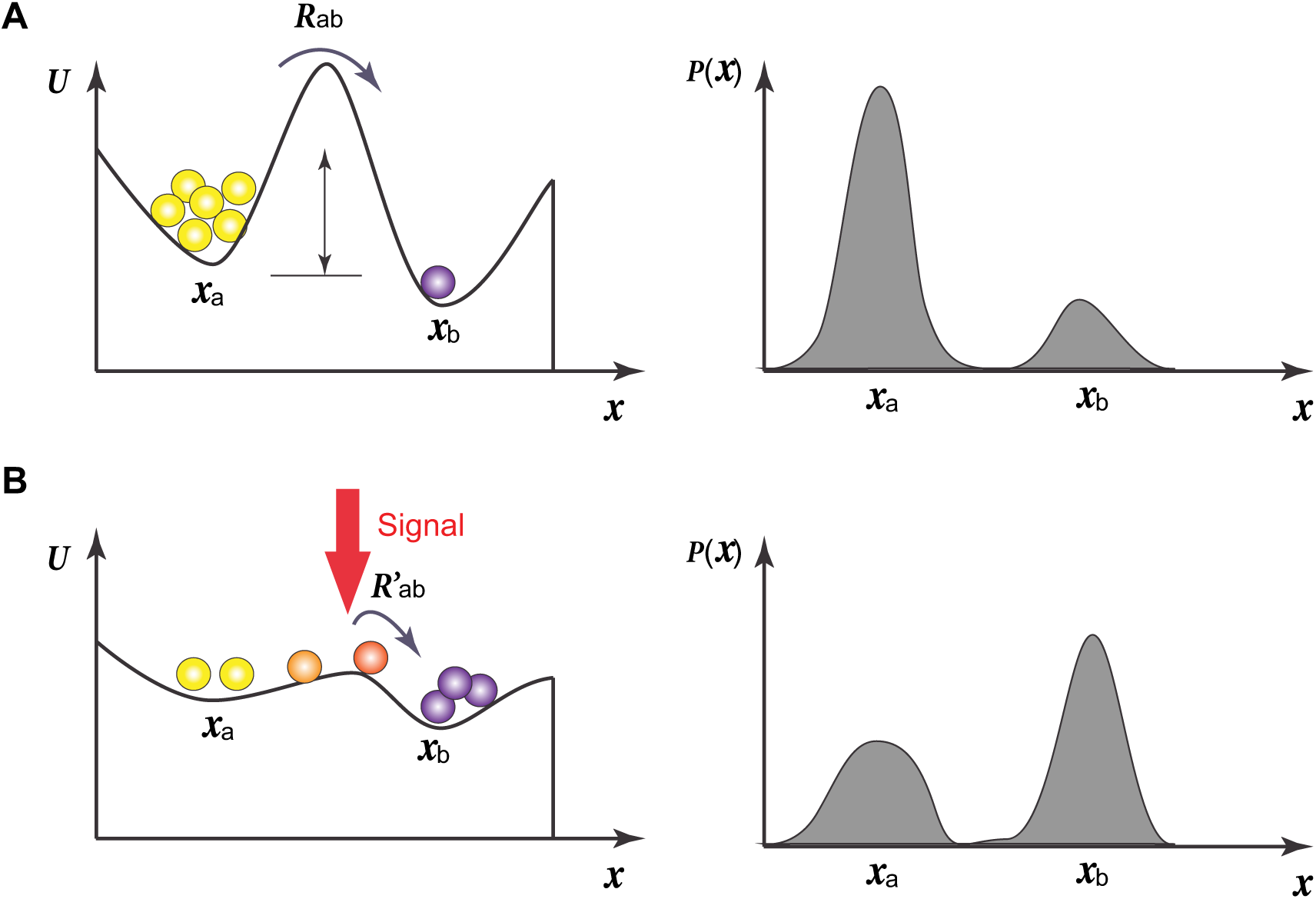
A cell state transition is driven by the quasi-potential (landscape) change induced by external signals. In (A) a bistable landscape exists with cells residing mainly in state *x*_*a*_ due to a high barrier to state *x*_*b*_. In (B) cells transit to *x*_*b*_, caused by the external signals that flatten the barrier.

However, single-cell technology and the measurement of high-dimensional states of many cells now provide a way to determine the *relative occupancy* probabilities of attractor states (density of clusters in state space) and *attractor transition rates* (at which a cell moves from one cluster to another). From these two measurements, we can phenomenologically obtain the landscape shape, such as relative sizes and depths of attractors, and the height of barriers between them, directly from single-cell states without knowledge of the specification of the dynamical system. The general idea is that the stochasticity of individual cells turns a cell population into a statistical ensemble that “reads out” the constrained state space as imposed by the gene regulatory network. For instance, from the cell density distribution in state space and at steady-state, we can define attractors. The transition rates between attractors can be revealed by sorting cells from one cluster and observing transitions to reconstitute another [6]. According to these transition rates, one can estimate their “relative stability” based on the theory of quasi-potential energies. A widely used intuitive approximation of the depth of an attractor is 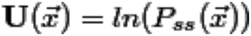, where 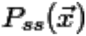 is the measured density of state 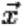 [66,67]. Note, however that a difference in this apparent potential is that this is not the source of the force that drives the state change: given the rate of state change and the quasi-potential 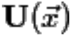, the driving force is not
simply [58]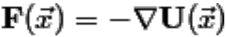[58]. Exact experimental determination of the transition probabilities between attractors would require longitudinal monitoring, in all relevant state dimensions, of cells undergoing such transitions – which is currently beyond reach. However, cell transition rates in lower dimensions can be measured [53] and have been modeled with a phenomenological bifurcation model [68].

Finally, a theory-based interpretation of snapshot data is based on the concept of ergodicity. In analogy to the ergodic hypothesis in statistical physics, a static picture of an ensemble of individual cells can inform us about the behavior of an individual over time – if the state changes of individuals are fast enough relative to the time of observation. The ergodic rate analysis (ERA) was used to estimate the rates of cell growth based on snapshots of a cell population [69], and population dynamics were deconvolved by tracking individual cells after cancer drug treatment [70].

## Conclusions

To understand the cell population substructures manifesting the constraints imposed by the underlying dynamical system which can be intuitively and formally depicted as a quasi-potential landscape, we need to go beyond descriptive computational data analysis and *ad hoc* interpretation and enter the still little charted terrain of theory-based interpretation. Here the most natural framework for understanding why the patterns in the data arise in the first place, is the theory of nonlinear stochastic dynamical systems [71]. In the near future we will see progress at all three fronts discussed in this article: In collecting data, the costs for profiling individual cells will drop drastically. For instance, Drop-seq for will make RNA-Seq with ten thousands of cells affordable, which is critical for a statistically robust evaluation of population substructures. Moreover, the combination of different profiling methods will enlarge the charted state space. At the front of correlating data points, we expect to see a consolidation. First-generation computational tools have served their purpose in introducing the intuition of single-cell resolution analysis of high-dimensional cell states, but lack a deeper understanding of the underlying regulatory system. At the front of connecting the dots, theory-based analysis will benefit from progress and sinking costs in data collection, which will permit the design of more complex experimental schemes with denser snapshots in order to test the theory. However, theoretical concepts, such as the quasi-potential landscape must be further developed and linked to data, and the abstract ideas need to be disseminated to a larger community of bioinformaticians.

## Acknowledgements

We thank Chris McGinnis (ISB, Seattle), and Michael Strasser, Alex Wolf, and Thomas Blasi (Munich) for comments on the manuscript, Philipp Angerer (Munich) for technical support, and the German Academic Exchange Service DAAD and Bavarian Research Alliance BayFOR for exchange funding. Research reported in this publication was supported by the German Science Foundation DFG (project ‘Inference of Differentiation Decision Times from Blood Stem Cell Genealogies’) and by the National Institutes of Health under award number R01GM987654. The content is solely the responsibility of the authors and does not necessarily represent the official views of the National Institutes of Health.

## Footnotes

* Molecular profiling of reprogramming dynamics of 36 markers in 250,000 mouse embryonic stem cells using mass cytometry.

* Gene expression profiling of nearly 4,000 mouse embryonic cells with blood forming potential using qRT-PCR. Computational analysis using diffustion maps and network reconstruction.

* Barcodes for individual molecules shown to allow for alleviating amplification noise and the assessment of transcript abundances.

* Introduction of Drop-seq, a droplet-microfluidic that allows handling and simultaneous sequencing of thousands of cells, applied to the identification of distinct cell populations in over 44,000 mouse retinal cells.

* Introduction of InDrop, similar to Drop-seq, applied to the characterization of mouse embryonic stem cell subpopulations.

* Computational approach to account for confounding factors in single cell gene expression data, applied to cell cycle induced variations.

* Application of the diffusion map concept to transcriptomics data of differentiation cells, including an overview of existing dimension reduction algorithms.

* Introduction of the Monocle method for pseudotemporal ordering of differentiating human muscle progenitor cells.

* Introduction of the Wanderlust algorithm that aligns single cells along a high-dimensional trajectory to reconstruct the developmental path of human white blood cell maturation.

* Quantitative time-lapse microscopy of transcriptional Nanog reporter in mouse embryonic stem cell reveals stochastic switching between gene expression states.

* Time-lapse data used to infer the Nanog DNA state from mRNA and protein kinetics in mouse embryonic stem cells.

* Landscape of gene expression variability determined by intercolony variance in transcript numbers of several pluripotency and lineage regulators with fluorescence in situ hybridisation (FISH), after 3-4 days of clonal growth, in traditional in-vitro and perturbed conditions.

* Analysis of cell state stability on the protein level, and transitions between them with thousands of single cells and millions of timepoints from quantitative time-lapse microscopy.

* Detailed theoretical construction of quasi-potential landscape based on Wentzell large deviation theory and its difference from U ~ ln P_SS_.

* Systematic description of the importance of landscapes in the context of evolution.

* Mathematical framework to deduce cell growth dynamics from flow cytometry snapshot data of single cells.

